# The cryptic origins of evolutionary novelty: 1,000-fold-faster trophic diversification rates without increased ecological opportunity or hybrid swarm

**DOI:** 10.1101/053140

**Authors:** Christopher H. Martin

## Abstract

Ecological opportunity is frequently proposed as the sole ingredient for adaptive radiation into novel niches. Alternatively, genome-wide hybridization resulting from ‘hybrid swarm’ may be the trigger. However, these hypotheses have been difficult to test due to the rarity of comparable control environments lacking adaptive radiations. Here I exploit such a pattern in microendemic radiations of Caribbean pupfishes. I show that a sympatric three-species radiation on San Salvador Island, Bahamas diversified 1,445 times faster than neighboring islands in jaw length due to evolution of a novel scale-eating adaptive zone from a generalist ancestral niche. I then sampled 22 generalist populations on seven neighboring islands and measured morphological diversity, stomach content diversity, dietary isotopic diversity, genetic diversity, lake/island areas, macroalgae richness, and Caribbean-wide patterns of gene flow. None of these standard metrics of ecological opportunity or gene flow were associated with adaptive radiation, except for slight increases in macroalgae richness. Thus, exceptional trophic diversification is highly localized despite myriad generalist populations in comparable environmental and genetic backgrounds. This study provides a strong counterexample to the ecological/hybrid-swarm theories of adaptive radiation and suggests that diversification of novel specialists on a sparse fitness landscape is constrained by more than ecological opportunity and gene flow.

## Introduction

Adaptive radiation is one of the most fundamental processes producing bursts of species diversification, trait divergence, and niche evolution across the tree of life (Simpson 1944; Schluter 2000). Ecological opportunity, an abundance of available resources in a new environment with few competitors, has long been viewed as the primary force driving diversification through divergent selection on niche use, mediated by competition and predation (Simpson 1944; Schluter 2000; Losos and Mahler 2010; Pfennig and Pfennig 2010). This relationship between new ecological space and divergent selection leading to reproductive isolation forms the foundation of the ecological theory of adaptive radiation (Schluter 2000; Losos and Mahler 2010) and ecological speciation theory (McKinnon et al. 2004; Rundle and Nosil 2005; Nosil 2012).

At microevolutionary scales, substantial observational and experimental evidence demonstrates that resource abundance drives ecological divergence (Schluter 2000; Parent and Crespi 2009; Nosil 2012). For example, parallel ecological diversification and speciation has been observed repeatedly in laboratory microcosms (Rainey and Travisano 1998; Elena and Lenski 2003; Kassen et al. 2004) and natural systems (Gillespie 2004; Langerhans et al. 2007; Losos 2009; Martin 2012) when populations are confronted with similar levels of ecological opportunity, constrained only by altered communities of predators or competitors (Vamosi 2003; Pfennig and Pfennig 2012). Holding lineage-specific factors and biotic communities constant, ecological divergence in response to resource abundance appears much more predictable than originally imagined (Bolnick and Lau 2008; Nosil et al. 2009; Langerhans 2010).

In contrast, it is still largely unknown what triggers major evolutionary novelties and large-scale adaptive radiation: why do only some lineages rapidly diversify and colonize novel ecological niches in response to resource abundance while others do not (Roderick and Gillespie 1998; Burns et al. 2002, Erwin 2015a)? Many macroevolutionary trends run counter to predictions of the ecological theory of adaptive radiation. First, contrary to Simpson’s paradigm (Simpson 1944), major ecological transitions to new adaptive zones are only weakly or not at all associated with the availability of ecological opportunity due to colonization of an isolated habitat, following mass extinction, or the evolution of a key innovation (Erwin 2015a; Harmon and Harrison 2015). For example, for nearly every adaptive radiation containing novel ecological specialists, there is a similar lineage which has failed to diversify in the exact same environment (Givnish et al. 1997; Roderick and Gillespie 1998; Arbogast et al. 2006; Losos 2009; Martin and Wainwright 2011, 2013c). Large-scale radiations often originate long before mass extinction events (Schuettpelz and Pryer 2009; Wilson et al. 2012) or the evolution of key innovations (Alfaro et al. 2009, Erwin 2015a). Moreover, most classic adaptive radiations do not exhibit an early burst of trait diversification, the rapid niche-filling response predicted by the ecological opportunity hypothesis (Harmon et al. 2010, Erwin 2015a; Harmon and Harrison 2015).

Second, the origins of adaptive radiation cannot be explained by the same factors that are associated with speciation and morphological diversification rates. Although there is a strong relationship between area and speciation rate (Losos and Schluter 2000; Kisel and Barraclough 2010), island biogeography breaks down when attempting to predict which lineages will radiate and where (Seehausen 2006; Wagner et al. 2012, 2014). Adaptive radiation also appears to defy ecological limits and frequently results in species assemblages exceeding the equilibrium species diversity expected from community assembly processes alone (Gillespie 2004; Gavrilets and Losos 2009; Martin and Genner 2009; Wagner et al. 2014). Thus, while ecological opportunity is sufficient to drive divergent selection among populations, its primary role in triggering macroevolutionary diversification is unclear (Erwin 2015a).

An emerging alternative to ecological opportunity for the origins of adaptive radiation is the hybrid swarm hypothesis, which proposes that an influx of genetic diversity and novel allelic combinations after hybridization of distinct lineages (inter- or intraspecific), possibly including segregating postzygotic intrinsic incompatibilities, may trigger rapid diversification (Seehausen 2004; Roy et al. 2015; Schumer et al. 2015). Hybridization plays a well-known role in the formation of single ecologically divergent species, even among homoploids (Rieseberg et al. 2003; Schumer et al. 2013). In contrast, although a growing number of studies identify substantial gene flow and adaptive introgression within adaptive radiations (Brawand et al. 2014; Lamichhaney et al. 2015; Malinsky et al. 2015; Martin et al. 2015, 2016; Stankowski and Streisfeld 2015), there is still no evidence that hybridization specifically triggered their diversification, as opposed to being pervasive throughout the history of these lineages (Servedio et al. 2013; Berner and Salzburger 2015; Kuhlwilm et al. 2016), and direct tests are needed. In order to directly test these hypotheses about the origins of adaptive radiation and novelty, we need to examine systems where parallel speciation is not the dominant feature and compare variation in ecological opportunity and hybridization across similar environments that differ in the presence of nascent adaptive radiations of specialists.

Here I exploit such a pattern of microendemic adaptive radiations and the rare evolution of novel trophic specialists in *Cyprinodon* pupfishes to test the roles of ecological opportunity and hybridization in triggering adaptive radiation and novelty. Pupfishes inhabit coastal areas and salt lakes across the entire Caribbean and Atlantic, from Massachusetts to Venezuela; however, despite the ubiquity of these populations and the abundant ecological opportunity present in most Caribbean salt lakes which lack predators and contain few competing fish species, nearly all populations are generalist omnivores, consuming algae and micro-invertebrates, and sympatric radiations of trophic specialists have evolved only twice: once in 10,000-year-old salt lakes on San Salvador Island, Bahamas and independently in the 8,000-year-old Laguna Chichancanab basin in the Yucatan (Martin and Wainwright 2011). In addition to a generalist algae-eating species present in all lakes (*C*. *variegatus*), the San Salvador radiation contains two rare specialist species coexisting with generalists in some lakes which have adapted to unique trophic niches using novel skeletal traits: a scale-eating pupfish with enlarged oral jaws [*C. desquamator* (Martin and Wainwright 2013a)] and a molluscivore pupfish with a nasal/maxillary protrusion which may provide jaw stabilization for crushing [*C. brontotheroides* (Martin and Wainwright 2013a)]. Similarly, the Chichancanab radiation of five species contains a generalist algae-eating species (*C. beltrani*) and at least two trophic specialists – a zooplanktivore (*C. simus*) and a piscivore (*C. maya*: (Humphries and Miller 1981; Humphries 1984; Stevenson 1992; Horstkotte and Strecker 2005). These trophic niches and skeletal phenotypes in each sympatric radiation are unique among all *Cyprinodon* species; furthermore, scale-eating is unique among over 1,500 species of Cyprinodontiform fishes (Martin and Wainwright 2011, 2013c). These unique traits are hypothesized to be novel, in the sense of a body part evolving quasi-independence or individuation through modification of gene regulatory networks (following Erwin 2015), and different scaling relationships with body size support this idea [Fig. S4; (Lencer et al. 2016)]. These traits are also innovations, in the sense of enabling access to new resources (following Liem 1980; Losos 2009; Blount et al. 2012a), but it is too early to determine if they are key innovations causing an increase in species diversification rate (Hunter 1998; Whittall and Hodges 2007).

Trophic specialists in both sympatric pupfish radiations co-occur in all lake habitats, but are largely reproductively isolated with low levels of hybridization (within-lake interspecific *F_st_* = 0.12-0.49: Strecker 2006a; Martin and Feinstein 2014). The San Salvador radiation is nested within many outgroup Caribbean populations of *C. variegatus*, strongly indicating that these specialists evolved from a generalist ancestor similar in morphology to *C. variegatus* (Martin and Wainwright 2011; Martin and Feinstein 2014). Similarly, the Chichancanab radiation is nested within outgroup populations of *C. artifrons* along the Yucatan coast, indicating that these specialists evolved from a generalist ancestor resembling *C. artifrons* (Martin and Wainwright 2011). Unfortunately, Chichancanab was colonized by two invasive fish species in the 1990’s which caused the extinction of at least two specialist species (Schmitter-Soto 1999; Strecker 2006b; Martin and Wainwright 2011). Thus, I focus on the San Salvador radiation in this study with occasional comparisons to initial descriptions of the Chichancanab environment before it was impacted by invasives [e.g. (Humphries and Miller 1981)].

In summary, two unusual features of Caribbean pupfishes provide an outstanding opportunity for investigating the causes of adaptive radiation and the rare evolution of ecological novelty: 1) major evolutionary novelties evolved within 10,000 years and are restricted to isolated locations, 2) each sympatric radiation is surrounded by a large number of comparable environments inhabited by a generalist lineage comparable to the putative ancestor where rapid adaptive diversification has not occurred. To pinpoint where major ecological novelties evolved and identify which ecological and genomic factors are associated with these transitions, I integrated population genomics, phylogenetic comparative methods, and a Caribbean-wide ecological, morphological, and genetic survey of pupfish populations. I first tested for the presence of exceptional trophic diversification and shifts to new adaptive zones by constructing a time-calibrated *Cyprinodon* phylogeny from 8,352 loci and measuring 28 functional traits in 22 pupfish populations on 7 islands (*n* = 493 individuals across all populations). I then tested whether ecological opportunity and hybridization were associated with the exceptional trait diversification rates detected on San Salvador. To test the role of ecological opportunity, I compared the best indicators of ecological opportunity in this system to the number of sympatric pupfish species within each lake: lake area, macroalgae species richness, and the genetic diversity, stomach content diversity, stable isotope diversity, and morphological diversity of generalist pupfish populations in each lake. To test the role of hybridization, I used genome-wide sampling of genetic markers to estimate whether the species on San Salvador showed evidence of hybrid swarm origins relative to neighboring islands using *f_4_* statistics, principal components analysis, species tree estimation, and Treemix population graphs.

## Methods

### Sampling

Twelve hypersaline lakes on San Salvador Island were sampled for both generalist and specialist pupfishes (if present), including four lakes containing only generalists, two containing generalists and molluscivores, two containing generalists and scale-eaters, and four containing generalists and both specialist species in July, 2011 (Fig. 1; Appendix S1). Seven hypersaline lakes containing generalist pupfish populations on five neighboring Bahamian Islands (Rum Cay, Cat, Long, Acklins, New Providence) and three lakes in the Dominican Republic (Laguna Bavaro, Laguna Oviedo, Etang Saumatre) were sampled between May – July, 2011 (sampling locations highlighted in Fig. 1 by brown arrows; Appendix S1). Additional outgroup *Cyprinodon* species were sampled from across the Caribbean and Atlantic, spanning the entire coastal range of *Cyprinodon* from Massachusetts to Venezuela (Appendix S1).

**Fig. 1.**
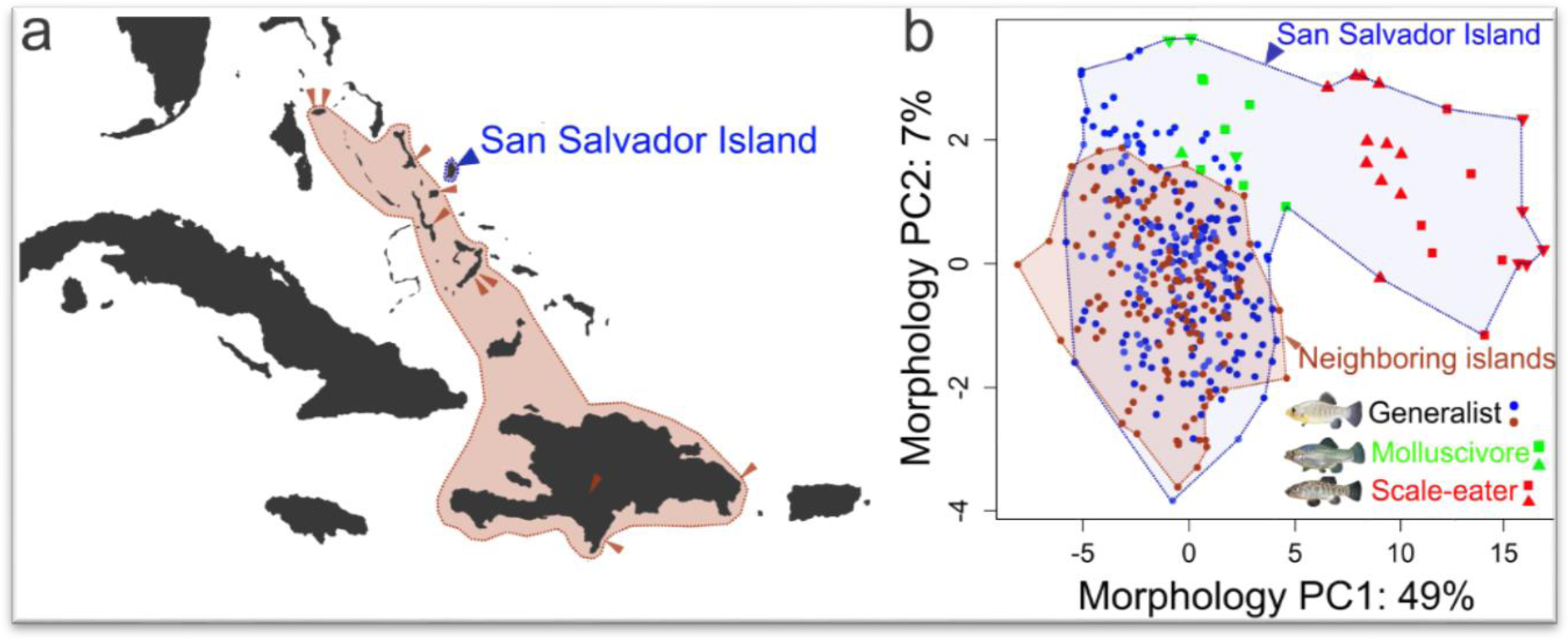
*a)* Sampling locations of generalist populations across the Caribbean (San Salvador lakes: *n* = 12; lakes on neighboring islands: *n* = 10). *b)* First two principal components of morphological diversity for 28 size-corrected skeletal traits measured on 493 alizarin-stained specimens (Fig. S3). Morphological diversity of generalists on San Salvador (●) is equal to the diversity of generalist morphology on neighboring islands (●). Diversity of all three San Salvador species (blue area) greatly exceeds trans-Caribbean diversity (brown area). Shapes indicate different specialist populations on San Salvador.

Neighboring lakes were comparable to those on San Salvador: hypersaline due to limited hydrological connectivity with the ocean dominated by mangroves in predominantly shallow (2 – 5 m) karst basins. Physiological parameters, including pH, salinity, alkalinity, and temperature are extremely variable within each lake due to tidal fluctuations and rainfall, but highly comparable across lakes due to their shared carbonate geology and water chemistry (Rothfus 2012). Lakes contained from 0 – 2 insectivorous fish species in addition to *Cyprinodon*: the Bahamian mosquitofish, *Gambusia hubbsi*, and the bighead silverside, *Atherinomorus stipes*, with the exception of both lakes on New Providence Island which contained invasive fishes such as *Oreochromis mossambicus, Xiphophorus maculatus*, and *Poecilia reticulata* and the three large lakes in the Dominican Republic, which contained cichlid, eleotrid, and American crocodile (*Crocodylus acutus*) predators and *Limia* spp. and *Gambusia* spp. competitors. Dominican lakes exhibited similar phenotypic and ecological diversity trends to those in the Bahamas and are included for comparison; removing these lakes did not qualitatively affect the results.

Between 9 – 48 specimens (mean = 21.6; sd = 8.8) were sampled from generalist populations in each lake by seine-net or hand net and euthanized in an overdose of buffered MS-222 (Finquel, Inc.) following animal care protocols approved by the University of California, Davis Institutional Animal Care and Use Committee (IACUC protocols #15908 and #17455).

### Genomic sequencing and bioinformatics

Between 1 - 6 wild-caught fish from each of 22 generalist populations, plus specialist *C. desquamator* and *C. brontotheroides* populations on San Salvador, and Caribbean-wide outgroup sampling spanning the entire Caribbean and Atlantic range of *Cyprinodon* (*n* = 112 individuals detailed in Appendix S1) were individually bar-coded and sequenced using the genotyping by sequencing RADseq protocol (Elshire et al. 2011), filtered reads were aligned to the *C. variegatus* reference assembly using bowtie2 (Langmead and Salzberg 2012), and genotypes were called using the Stacks pipeline (Catchen et al. 2013), exporting loci with a minimum depth of 10 reads genotyped in >50% of individuals following the approach of previous RADseq studies (Martin and Feinstein 2014; Martin et al. 2015, 2016; Martin et al. in review). Respectively, 601 million 150-bp and 356 million 100-bp raw reads were sequenced and 651 million reads uniquely aligned to the *C. variegatus* assembly. Further details are provided in the supplemental methods.

### Testing for exceptional diversification rates: phylogenetic and comparative analyses

I first used BEAST (Drummond and Rambaut 2007) to estimate a time-calibrated phylogeny for Caribbean *Cyprinodon* populations from 8,352 concatenated RADseq loci (following Martin et al. 2015), detailed in the supplemental methods and presented in Fig. S1. I then used two complementary approaches to assess the distribution of trait diversification rates across this phylogeny. First, I used reversible-jump MCMC sampling of multi-rate Brownian motion models for each trait on the maximum likelihood phylogeny (Eastman et al. 2011). Nearly all nodes in the time-calibrated tree were resolved with posterior probabilities of 1 (Fig. S1); therefore, phylogenetic uncertainty was not accounted for in these analyses and only a single tree was used. Reversible-jump sampling allows for ‘jumps’ among models with varying numbers of parameters in addition to varying the values of these parameters while sampling likelihood space, enabling the MCMC sampler to explore shifts among different diversification rate regimes on different parts of the phylogeny without *a priori* specification of these shifts as required in earlier methods (O’Meara et al. 2006). This approach also naturally results in model-averaged parameter estimates (Burnham and Anderson 2002). For each residual trait measured in 29 taxa (see below: *Morphological diversity*), I used the *auteur* suite of methods (Eastman et al. 2011), part of the geiger2 package (Harmon et al. 2008) in R (R Core Team 2015), to run rjMCMC chains and estimate the placement and magnitude of shifts in diversification rate across the tree. Each chain was run for 10,000 generations, sampling every 100 steps from a relaxed Brownian motion model allowing for the possibility of instantaneous jumps between rate regimes (*type* = “jump-rbm”). The first 50% of each chain was discarded as burn-in and shifts in rate regimes were visualized on the phylogeny for each trait. Chains were run multiple times for each trait to assess convergence. Median diversification rates and their placement on the phylogeny for rapidly-diversifying traits was generally robust across runs; however, this is primarily an exploratory technique for visualizing the highest densities of major trait diversification shifts and no explicit model comparisons were performed (e.g. Santini et al. 2013).

Second, I used reversible-jump MCMC sampling of the more generalized OrnsteinUhlenbeck (OU) model of trait diversification. In addition to a Brownian motion parameter (σ^2^) modeling diffusion rate, the OU model includes the elastic pull of a stabilizing selection regime acting on a trait with two additional parameters: α, the strength of selection on the trait, and θ, the location of the fitness optimum (Hansen 1997; Martins and Hansen 1997; Butler and King 2004). Stronger selection on a trait as it nears its fitness optimum constrains the stochastic diffusion rate of σ^2^ over time, erasing the phylogenetic signal of a trait (Hansen et al. 2008). Although a single fitness optimum can be fit to an entire clade, the ecological theory of adaptive radiation predicts that multiple fitness peaks will drive ecological divergence and speciation (Schluter 2000), indicating that multi-optima OU models are needed to model adaptive radiation. Indeed, multiple fitness peaks on a complex fitness landscape were directly measured in this system (Martin and Wainwright 2013c), demonstrating that fitness peaks were non-Gaussian and connected by varying depths of fitness valleys and ridges on a single adaptive landscape for the entire three-species radiation, in contrast to instantaneous transitions among single-optimum Gaussian fitness regimes along branches specified by the multi-rate OU. Nonetheless, OU models may provide a rough approximation of stabilizing selection acting on different phenotypic optima over evolutionary time and enable estimation of the temporal dynamics of adaptive regime shifts across the *Cyprinodon* phylogeny.

To specifically explore jaw length, the trait with the largest shift in diversification rate on San Salvador, I used bayou to conduct reversible-jump MCMC sampling of multi-rate OU models (Uyeda and Harmon 2014) in R. I first specified an uninformative prior on the OU models, using half-cauchy priors for α and σ^2^, a normal prior for θ, a conditional Poisson distribution for the number of shifts between selective regimes (lambda = 15, max = 200), and a maximum of one shift per branch. I also explored more restrictive priors for the number of shifts, but found convergence across runs to be less stable. I ran two chains of 1,000,000 generations each, sampling every 100 steps, and combined chains after discarding the first 50% as burnin-in. Convergence was assessed using Gelman and Rubin’s R statistic. Repeated runs indicated that convergence was quickly reached and all runs indicated a robust shift to a new adaptive regime for scale-eaters. Results were plotted as a ‘traitgram’ (Ackerly 2009) using bayou’s built-in plotting function (Uyeda and Harmon 2014).

### Testing the ecological opportunity hypothesis: morphological, dietary, and genetic diversity

To test the hypothesis that ecological opportunity is associated with exceptional diversification rates on San Salvador Island, I used linear models to measure the effects of morphological diversity, dietary diversity, genetic diversity, lake/island areas (from the Google Maps Area Calculator Tool: daftlogic.com), and macroalgae species richness on the number of pupfish species coexisting within a lake (ranging from 1-3 species; Tables 1–2). Multiple regression models were compared using stepwise addition and removal of predictors with the stepAIC function in the MASS package in R (Venables and Ripley 2013). The fit of significant one-way models was compared to full models including all interaction terms. Estimation of predictor variables is described below.

**Table 1.** Association between island/lake area, ecological diversity, genetic diversity, and morphological diversity with the number of sympatric *Cyprinodon* species coexisting within a lake (*n* = 1 – 3) across 22 lake populations on 7 islands (Fig. 4). Except for island area, all linear regressions treated each lake population as independent replicates; however, note that degrees of freedom in these tests may be inflated by the additional covariance among populations due to their varying degrees of shared history (Felsenstein 1985; Revell 2009). The only significant positive correlation is highlighted in bold.

| variable | correlation with *Cyprinodon* spp. | $r^2$ | $P$ |
| --- | --- | --- | --- |
| $\log_{10}$ island area | *negative* | 0.695 | 0.003 |
| $\log_{10}$ lake area | - | 0.006 | 0.654 |
| genetic diversity | - | 0.030 | 0.284 |
| **macroalgae richness** | **positive** | **0.416** | **0.002** |
| stomach content diversity | - | 0.003 | 0.856 |
| $\delta^{13}\text{C}$ | - | 0.001 | 0.903 |
| $\delta^{15}\text{N}$ | - | 0.012 | 0.646 |
| PC1 variance | - | 0.131 | 0.107 |
| PC2 variance | - | 0.044 | 0.360 |
| LD1 variance | - | 0.017 | 0.571 |
| LD2 variance | - | 0.025 | 0.496 |

**Table 2.** Models incorporating the effects of lake area, ecological diversity, genetic diversity (π), and morphological diversity on the number of sympatric *Cyprinodon* species coexisting within a lake (spp. = 1 – 3). Linear regression models treated the ecology of each lake as independent replicates; however, note that degrees of freedom for comparisons of generalist traits may be inflated by the additional covariance among populations due to their varying degrees of shared history (Felsenstein 1985; Revell 2009). Significant models are highlighted in bold and include presentation of significance, effect sizes, and SE for each term.

| **model** | ***df*** | ***adj.r<sup>2</sup>*** | ***P*** |
| --- | --- | --- | --- |
| spp. ~ log(lake area)+$\pi$+macroalgae richness+stomach content diversity+$\delta^{13}\text{C}$+$\delta^{15}\text{N}$+PC1 variance+PC2 variance+LD1 variance+LD2 variance | 10 | 0.10 | 0.546 |
| spp. ~ log(lake area)+$\pi$+macroalgae richness+PC1 variance+PC2 variance+LD1 variance+LD2 variance | 16 | 0.32 | 0.216 |
| **spp. ~ macroalgae richness+PC1 variance+ LD1 variance** | **16** | **0.53** | **0.005** |
| **macroalgae richness:** 0.223 ± 0.07 | | | **0.009** |
| PC1 variance: 0.176 ± 0.12 | | | 0.162 |
| LD1 variance: 1.054 ± 0.50 | | | 0.057 |
| spp. ~ macroalgae richness*PC1 variance*LD1 variance | 16 | 0.49 | 0.506 |

#### Morphological diversity

Multiple individuals from 21 lake populations (*n* = 493 individuals total; 9 – 48 samples per population; mean = 21.6; sd = 8.8) were cleared and alizarin-stained for measurement of 28 functional skeletal traits (Figs. S2-S4). For each population, adult specimens were cleared and double-stained with alizarin and alcian blue in order to visualize skeletal morphology (Dingerkus and Uhler 1977). The skull of each specimen was photographed on both lateral sides with jaw adducted for a clear view of the quadroarticular region, framing only the head and pectoral girdle for maximum resolution of smaller features (Fig. S3). Specimens were photographed and measured on both lateral sides of the head and the mean was used to reduce measurement error. Thirty-two landmarks (Fig. S3) were digitized using tpsdig2 software (Rohlf 2001) and converted to 29 linear distances (Table S1) describing functional traits and the most divergent traits among the three San Salvador species.

After removing or remeasuring outlier measurements in the dataset, log-transformed linear distances were regressed against log-transformed head size of *C. variegatus* individuals as an index of overall size (Fig. S4). Size-corrected residuals were used for surveys of morphological diversity across generalist populations. Residuals from a separate linear regression calculated from the mean of each population were used for phylogenetic comparative analyses so that each taxon was weighted equally for the size-correction procedure.

#### Dietary diversity

For a subset of the generalist individuals sampled from each population, dietary diversity was estimated from stomach content analyses (*n* = 359 individuals total) and stable isotope analyses of δ13C and δ15N (*n* = 487 individuals total; Fig. S5). These complementary approaches reflect short-term fine-grain and long-term coarse-grain estimates of dietary diversity, respectively (Post 2002; Layman 2007). Stomach items were separated into broad taxonomic categories (e.g. macroalgae, seagrass, polychaete, ostracod, gastropod) and total surface areas of each component were estimated under 10-50x magnification using a Sedgwick-rafter cell [following (Martin and Wainwright 2013c)]. Dietary diversity within each population was estimated from Simpson’s inverse diversity index, the probability that two randomly sampled items do not belong to the same category (DeJong 1975). For stable isotope analyses, dried muscle samples from the caudal peduncle region of each fish were analyzed for natural abundances of δ13C and δ15N at the UC Davis Stable Isotope Facility on a PDZ Europa ANCA-GSL elemental analyzer interfaced to a PDZ Europa 20-20 isotope ratio mass spectrometer.

#### Genetic diversity

Genetic diversity across lake populations of generalist *Cyprinodon* was estimated by exporting 170 loci genotyped completely in 22 populations to avoid any biases introduced by missing data and using Stacks to calculate π, the average number of pairwise differences between sequences drawn at random within the population (Catchen et al. 2013). Estimates of genetic diversity were similar and qualitatively unchanged when running analyses with more permissive filtering criteria, at the island-level, and without genome alignment.

### Testing the hybrid swarm hypothesis: major axes of genetic variation and introgression

#### Principal components analysis

The main prediction of the hybrid swarm hypothesis is that hybridization will facilitate the evolution of ecological specialists. To test for this pattern, I first estimated principal components of genetic variation across the Caribbean, excluding San Salvador populations. 11,706 SNPs genotyped in at least 50% of individuals with a minimum depth of 10 aligned reads were exported for principal component analyses of genetic variance using probabilistic pca in the Bioconductor pcaMethods package (Stacklies et al. 2007) in R. Only 1 SNP per locus was exported to reduce linkage disequilibrium within this dataset. I then projected all three San Salvador species (*n* = 75 individuals) onto the first two principal components of Caribbean-wide genetic variation to assess whether specialists shared a greater proportion of their ancestry with any of the outgroups sampled. This approach has previously been used to visualize patterns of shared ancestry within a focal population while avoiding biases introduced by uneven sampling among groups inherent to principal components analysis (e.g. see discussion in McVean 2009; Lazaridis et al. 2014; Martin et al. 2015).

#### Species tree inference

Concatenation approaches may be consistently misleading estimators of the species tree, even with large amounts of data (Degnan and Rosenberg 2006, 2009; Kubatko and Degnan 2007; Heled and Drummond 2010; Liu et al. 2015). To estimate the species tree for Caribbean pupfishes, I used SNAPP, which integrates over all sampled gene trees (Bryant et al. 2012). To limit the computational demands of this analysis, I restricted the dataset to 1,534 SNPs genotyped completely in 21 focal Caribbean populations (Appendix S1: *n* = 70 individuals; mean = 3.3 per population) and pooled closely related saline lake populations on Crooked/Acklins Islands, New Providence Island, and Long Island were each pooled. Only 1 SNP was sampled per RAD locus to reduce linkage disequilibrium. Using BEAST2 (v. 2.2.0; Bouckaert et al. 2014) with the SNAPP plug-in (Bryant et al. 2012), two chains were run for 150,000 and 275,000 generations, respectively, and converged after 60,000 and 100,000 generations of burn-in, assessed using Tracer (v. 1.6; Drummond and Rambaut 2007). Due to slow run times on an 8-core Pentium i7 machine (1 million generations every 8,000 hours or 333 days), large effective sample sizes of parameters were not obtainable, which ranged from 3 – 170, affecting several theta parameters. After discarding burn-in, trees were sampled every 100 generations from both runs and visualized using Densitree in BEAST2 (Bouckaert et al. 2014).

#### Tests of asymmetric gene flow using f4 statistics

Species trees may still be an inadequate model to understand population history if secondary or continuous gene flow is present (Pickrell and Pritchard 2012). I used *f_4_* statistics to test for additional gene flow between branches within four-population maximum likelihood trees containing the two specialist species relative to Caribbean outgroups of the form: ((*C. brontotheroides, C. desquamator)*; (Caribbean outgroup 1, Caribbean outgroup 2)). This statistic measures asymmetry in allele frequency correlations between the (A,B) and (C,D) clades on the four-population tree ((A,B);(C,D)) and is expected to be zero in the absence of gene flow with only incomplete lineage sorting (but may be sensitive to ancestral population structure: Reich et al. 2009; Durand et al. 2011). Similar to the ABBA-BABA/D-statistic, the *f_4_* statistic tests for evidence of secondary gene flow in focal taxa, but does not require a rooted four-taxon tree (Martin et al. 2015). Treemix software [v. 1.12; (Pickrell and Pritchard 2012)] was used to calculate *f_4_* statistics from 4,213 SNPs genotyped in all focal Caribbean populations (pooled as described for SNAPP analyses) and sampled once per RAD locus.

#### Treemix population graphs

To visualize secondary gene flow among Caribbean populations, I used Treemix [v. 1.12; (Pickrell and Pritchard 2012)] to estimate maximum likelihood trees with varying numbers of migration events connecting populations after their divergence, forming graphs of interconnected populations. The number of migration events was estimated by comparing the log likelihood of graph models following global realignment and jackknife estimation in windows of size 1 using the 4,213 SNP dataset (following Martin and Feinstein 2014). To estimate the number of migration events connecting populations, I used an approach similar to Evanno et al. (2005) and compared the rate of change of the log likelihood with the addition of each migration event.

## Results

### Exceptional trophic diversification rates within San Salvador Island pupfishes

I found strong evidence for exceptional rates of trophic diversification and shifts to new adaptive zones localized to San Salvador Island, Bahamas (Fig. 2) despite extensive sampling of saline lakes with identically depauperate fish communities on 5 neighboring Bahamian islands and the three largest Caribbean saline lakes found in the Dominican Republic (Fig. 1). Sampling from the posterior distribution of multi-rate Brownian motion models indicated that the fastest trait diversification rates occurred for jaw length on the internal branch leading to all three populations of the scale-eating pupfish, *C. desquamator*, endemic to San Salvador (Fig. 2a, Table S1). These analyses must be interpreted with caution due to the small size of our phylogeny and violation of *auteur*’s assumption of a bifurcating phylogeny (see Fig. 5); however, the relative rate difference observed was substantially larger than the largest effect size used in *auteur* simulations and larger effect sizes increase precision in the inferred placement of true rate shifts in this model (see Fig. 2 in Eastman et al. 2011). Nonetheless, comparative methods for population networks are needed.

**Fig. 2.**
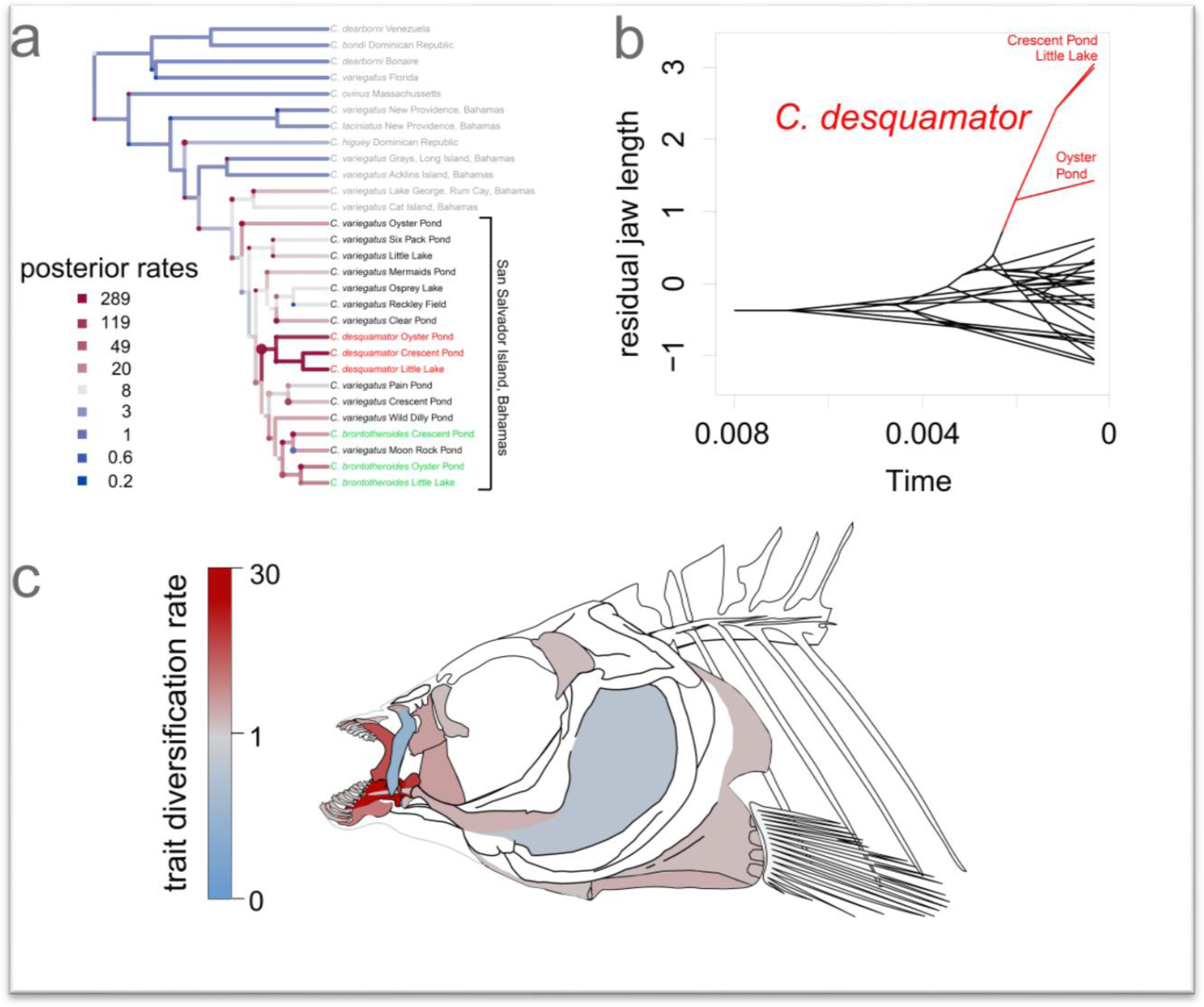
***a)* Shifts in jaw diversification rate** painted along each branch of the phylogeny estimated from mean residual jaw size in each population. Heat colors along each branch indicate median rates estimated from the posterior distribution of models and the size of circles at each node indicate the probability of a shift to a different rate regime. There was a 49% posterior probability of a shift to the highest jaw diversification rate at the root node of all three scale-eating pupfish populations, which was 30.8 times faster than median background diversification rates (Table S1) and **1,445 times faster than the slowest background rate** (legend in panel *a*: 289/0.2) on most neighboring islands. ***b)* Traitgram** illustrating distribution of mean residual jaw lengths across *Cyprinodon*. Red branches indicate the estimated phylogenetic position of a new adaptive regime corresponding to all three scale-eating populations (*C. desquamator*) with greatly enlarged jaws relative to the ancestral adaptive regime for generalist pupfish indicated by the black branches. ***e) Cyprinodon* anatomy-gram** illustrating median trait diversification rates on San Salvador relative to background rates on neighboring islands (modified from original heat map diagram in Martin and Wainwright 2011). Warm/cool colors indicate faster/slower rates, respectively. Representative skeletal regions are highlighted for a selection of linear traits presented in Table S1.

Accelerated trait diversification rates were most pronounced for jaw length and the width of the articular coronoid process (Fig. 2c, Table S1), which forms the base of the jaw closing lever and reflects the biomechanical tradeoff between fast scale-eating strikes and powerful shell-crushing force in the San Salvador specialists. Only two of the 28 traits measured showed decelerated trait diversification rates on San Salvador relative to other islands (Fig. 2c, Table S1). Sampling from the generalized Ornstein-Uhlenbeck model of trait evolution strongly supported a shift to a new adaptive regime for scale-eating (Fig. 2b). The highest posterior probability of a shift to a new fitness optimum was observed for jaw length along the internal branch leading to scale-eating pupfish (posterior probability = 0.25: Fig 2b). This new scale-eating optimum jaw size was nearly three standard deviations away (in mean standardized units) from the jaw length optimum estimated for generalist pupfish, indicating a major phenotypic transition (Fig. 2b).

### Increased ecological opportunity is not associated with adaptive radiation and trophic novelty

The ecological theory of adaptive radiation predicts that such exceptional rates of morphological, ecological, and sympatric species diversification on San Salvador should be associated with increased ecological opportunity. In contrast, I found no association between the presence of ecological specialist pupfishes and variation in nearly every standard index of ecological opportunity. First, the predicted island biogeographic relationship between island size and speciation rate (Losos and Schluter 2000; Kisel and Barraclough 2010; Wagner et al. 2014) does not hold up in this system. There was no correlation between lake area and the number of sympatric pupfish species and a negative correlation between island area and sympatric pupfish species (Table 1, Fig. 3a-b). Second, despite theoretical predictions (Weinreich and Chao 2005), genetic diversity within lake populations, an indicator of effective population size, was not associated with the number of sympatric pupfish species (Fig. 3c). Comparable levels of genetic diversity were also observed in the Chichancanab radiation (CHM unpublished data). Instead, low genetic differentiation among Caribbean islands is consistent with previous demonstrations of pervasive gene flow among islands and may continually renew genetic diversity within saline lakes (Martin and Feinstein 2014; see also Martin et al. 2016). Third, in contrast to many similar studies demonstrating a positive correlation between ecological opportunity and intraspecific or interspecific morphological and dietary diversity (Losos and Schluter 2000; Burns et al. 2002; Parent and Crespi 2009), an extensive survey of generalist populations across San Salvador and neighboring islands provided no evidence of increased ecological or phenotypic diversity in San Salvador lakes supporting specialists (Tables 1–2, Figs. 4, S2, S5). The diets of generalist pupfish should most directly reflect the availability of accessible resources within an environment, the best possible measurement of ecological opportunity. Furthermore, in contrast to expectations of character displacement (Pfennig and Pfennig 2012), there was no significant difference in dietary or morphological diversity between generalist populations on San Salvador with and without specialist pupfish species present (Table S2), suggesting that San Salvador generalist populations provide a good ecological baseline for comparison to other islands. There was no association between lakes supporting specialists and stomach content diversity, nitrogen isotopic diversity [indicating relative trophic position (Post 2002)], carbon isotopic diversity (indicating dietary carbon sources), or morphological diversity of generalist populations (estimated from principal component axes and discriminant axes separating the three San Salvador species based on 28 skeletal traits: Figs. 4, S2, S5; Tables 1–2).

**Fig. 3.**
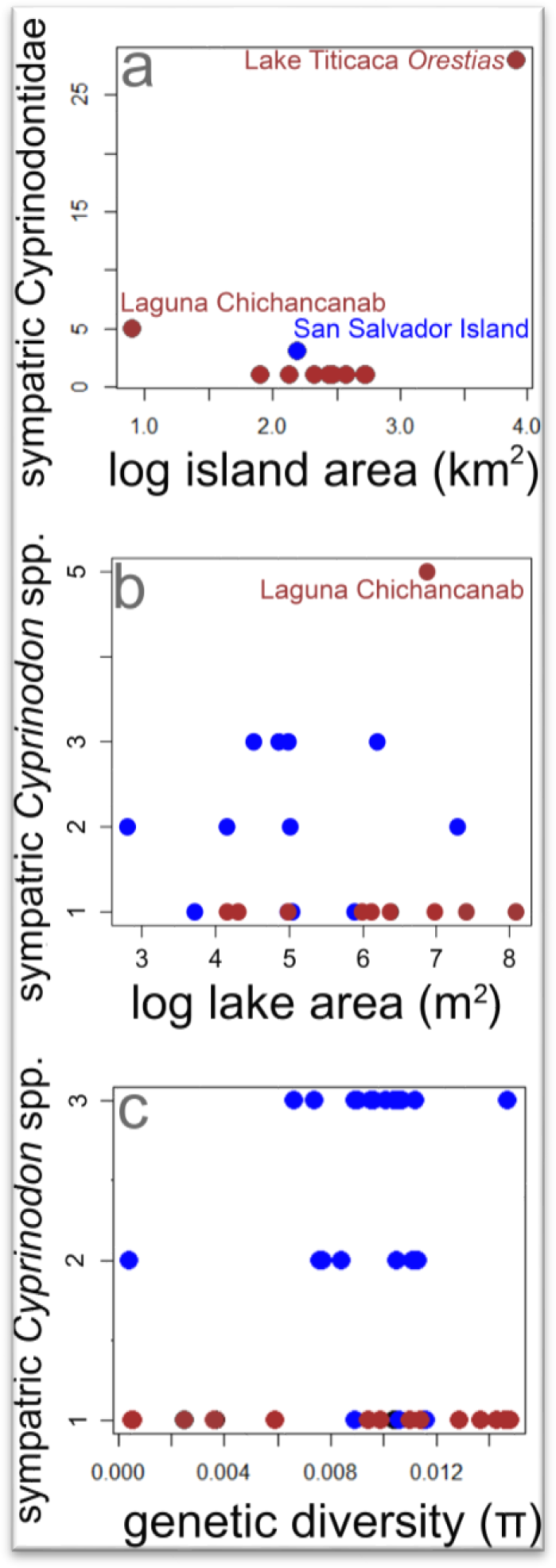
In contrast to the expected speciation-area relationship, there was no correlation between *a)* island area, *b)* lake area, or *c)* genetic diversity and the number of endemic *Cyprinodon* species (Tables 1–2). San Salvador populations are highlighted in blue, all other populations on neighboring islands in brown. The predicted speciation-area relationship was only supported at a much larger scale by including the distantly-related radiation of *Orestias* pupfishes in Lake Titicaca [log lake area: *r^2^* = 0.24, *P* = 0.003; log island area: *r^2^* = 0.364, *P* = 0.050; (Vila et al. 2013)].

**Fig. 4.**
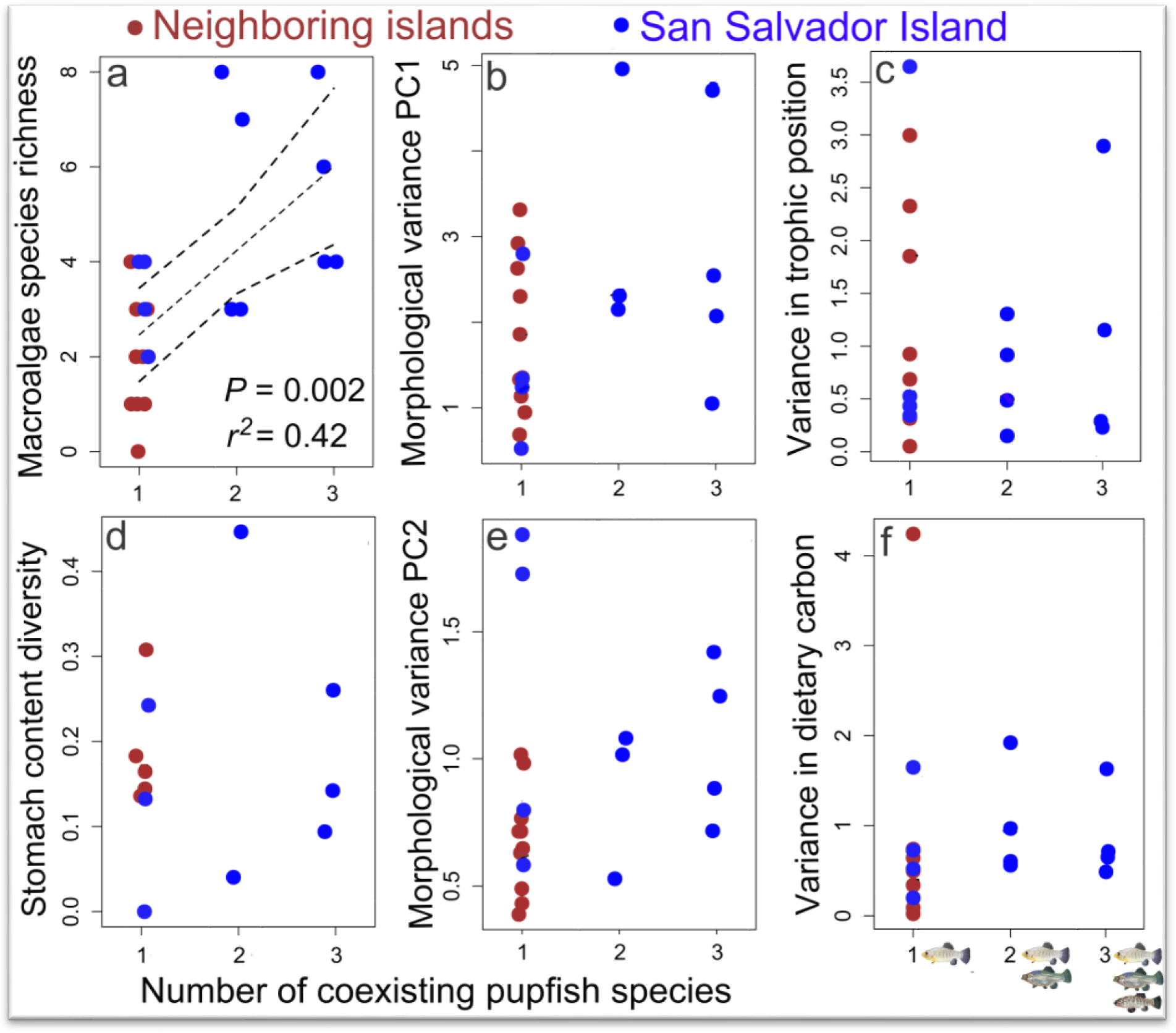
*a)* Macroalgae species richness was significantly correlated with the number of coexisting pupfish species within a lake (dashed lines indicate best-fit regression line and 95% confidence interval; Table 1). All other ecological and morphological variables were not correlated with the number of coexisting pupfish species: *b,e)* morphological diversity on the first two principal component axes (*n* = 493 individuals, 21 populations), *d)* stomach content diversity (Simpson’s index: *n* = 359 individuals, 13 populations), and *c,f)* dietary isotopic diversity (δ15N: trophic position and δ13C: dietary carbon) of generalist populations in Caribbean salt lakes (*n* = 487 individuals, 20 populations). Only generalist populations were compared among all sites.

The only significant effect uncovered among all univariate and multivariate models was the greater species richness of macroalgae communities within some lakes on San Salvador supporting multiple pupfish species (Fig. 4a, Tables 1–2). However, these additional macroalgae species (e.g. *Ulva fluxuosa, Valonia ventricosa*) made up less than 0.1% of total plant biomass and these lakes were still dominated by three macroalgae species (*Batophora oerstedii, Cladophora crispata*, and *Acetabularia crenulata*) and one marine angiosperm (*Ruppia maritima*) found throughout the Bahamas (Godfrey 1994). Furthermore, only half the lakes containing specialists had higher macroalgae richness (Fig. 4a). It is also notable that only a single macroalgae species (*Chara* sp.) occurs in Laguna Chichancanab (Humphries and Miller 1981), demonstrating that a diverse macroalgae community is not necessary for adaptive radiation. Macroalgae richness also exhibited no significant interactions with dietary diversity or lake size (Table 2) and the only other variable with a marginal effect on sympatric species number was morphological diversity along discriminate axis 1 (LD1), suggestive of slightly increased variation in jaw length (which loads heavily on this axis) in populations with specialists, most likely due to elevated within-lake introgression (demonstrated in Martin and Feinstein 2014).

### Increased gene flow is not associated with adaptive radiation and trophic novelty

Alternatively, the hybrid swarm theory of adaptive radiation predicts that adaptive radiation should be uniquely tied to admixed populations receiving a large influx of gene flow from highly divergent surrounding lineages (Seehausen 2004). This hypothesis must be evaluated with caution because gene flow is pervasive during adaptive radiation, but is often not unique to the radiating lineage (Abbott et al. 2013; Seehausen et al. 2014; Berner and Salzburger 2015). Indeed, there was substantial evidence for secondary gene flow with the specialist populations on San Salvador based on asymmetric allele frequency correlations with outgroups (significant *f_4_* statistics in 20 out of 55 four-population tests: Table S3), but also with generalist populations on neighboring islands that failed to radiate (Fig. 5). This can visualized as the likelihood of secondary gene flow connecting island populations in Treemix population graphs (Fig. 5c-d). Support for highly interconnected population graphs did not subside until reaching 20 connections (Fig. 5c-d), supporting widespread gene flow among Caribbean islands in violation of a bifurcating branching structure assumed by phylogenetic models (in contrast to Figs. 2a, 5b, S1). Importantly, the inferred directions of gene flow events among Caribbean islands crisscrossed the entire archipelago, rather than forming an epicenter at San Salvador, in contrast to the predictions of the hybrid swarm hypothesis (Fig. 5c-d).

**Fig. 5.**
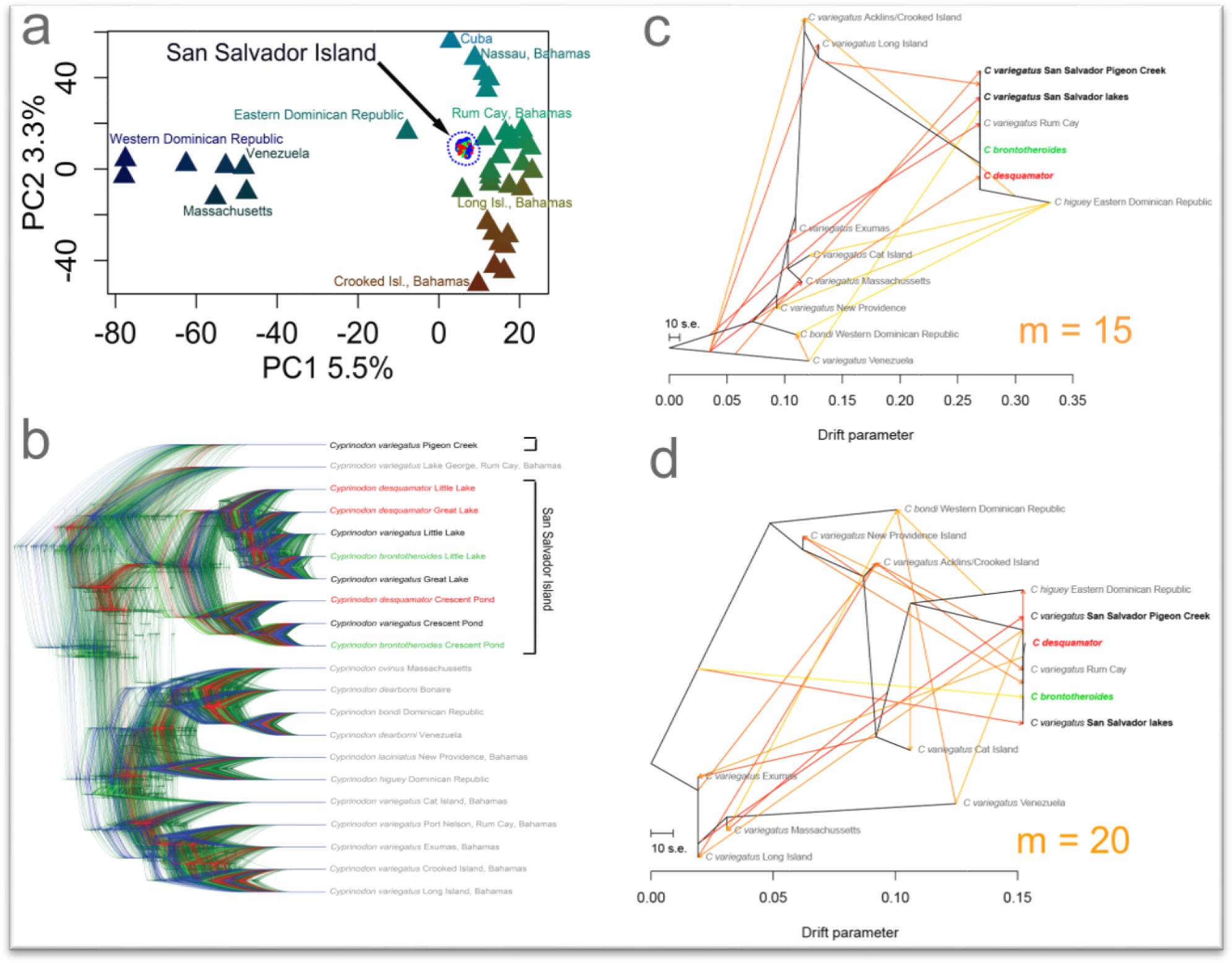
***a)*** Genetic variation within San Salvador populations relative to the genetic variation across the Caribbean for 11,706 SNPs. Individuals of all three species on San Salvador (●generalist *n* = 33; ●molluscivore *n* = 24; ●scale-eater *n* = 19) were projected onto the first two principal component axes of Caribbean-wide genetic variation (excluding San Salvador Island). Outgroup populations (▲) are colored according to their values along each principal component axis. ***b)* Distribution of gene tree topologies** estimated from 1,534 RAD loci (1 SNP per locus) using SNAPP plugin in BEAST2 (Bryant et al. 2012). ***c-d)*** Treemix population graphs (Pickrell and Pritchard 2012) illustrating gene flow among Caribbean populations based on stepwise fitting of migration events (shown as colored arrows, heat color indicates amount of gene flow) to a maximum likelihood phylogeny for 4,213 SNPs genotyped in all populations. Graphs are illustrated with 15 and 20 migration events (*c* and *d*, respectively). Following the approach of Evanno for choosing the number of genetic clusters in a dataset (Evanno et al. 2005), the rate of change of the likelihood began to decline as the number of migration events approached 20 (*m* = 10: lnL = 97; *m* = 15: lnL = 350; *m* = 20: lnL = 552; *m* = 25: lnL = 550).

Projections of San Salvador populations onto the first two principal components of genetic variation across the Caribbean indicated a pattern of continuous variation across islands, often diverging from geographic proximity (Fig. 5a). For example, most introgression into San Salvador specialists came from *C. higuey* in the Eastern Dominican Republic (Table S3; Fig. 5), which was only distantly related to western Dominican Republic populations.

Species tree analysis illustrated the diversity of majority-consensus phylogenies across the sampled RAD loci, including many different topologies for the sister group relationships among San Salvador specialist populations and outgroups on neighboring islands (Fig. 5b; however, note that a bifurcating tree is an inadequate model of gene flow in this system). For example, there was consistent support for intra-lake clustering of all three species in Crescent Pond, consistent with genome-wide introgression and islands of genetic differentiation (Martin and Feinstein 2014), but diverse topologies supported sister relationships between the San Salvador radiation and a coastal population of *C. variegatus* (Pigeon Creek, San Salvador Island), the closest island Rum Cay, or more distant islands (Fig. 5b).

## Discussion

Overall, these results strongly indicate that an exceptional increase in trait diversification rates and a major ecological transition from an ancestral generalist diet of algae and microinvertebrates to a new adaptive zone for scale-eating occurred only on San Salvador Island, Bahamas. This might reflect the extreme functional demands of scale-eating (Sazima 1983; Janovetz 2005) and the ecological novelty of this niche within pupfishes, which is separated by 168 million years from other extant scale-eating fishes (Martin and Wainwright 2013c). These estimates of relative trait diversification rates are similar to a previous species-level study across *Cyprinodon* using a mitochondrial phylogeny (Martin and Wainwright 2011) and substantially exceed relative diversification rates for the same jaw traits measured in classic examples of adaptive radiation in fishes, including Malawi cichlids [up to 9-fold faster than New World cichlids (Hulsey et al. 2010)] and coral reef fishes [2-fold faster than non-reef (Price et al. 2011)]. Indeed, the observed relative rates of trait diversification are among the fastest ever reported (Barkman et al. 2008; Ackerly 2009; Harmon et al. 2010).

In contrast to the large literature documenting ecological drivers of speciation (Schluter 2000; McKinnon et al. 2004; Rundle and Nosil 2005; Nosil et al. 2009; Parent and Crespi 2009; Nosil 2012), variation in ecological opportunity among depauperate saline lake environments does not appear sufficient to explain the rare occurrence of adaptive radiation in pupfishes. The young age and homogeneity of carbonate saline lakes across the Bahamas suggest that similar levels of ecological opportunity also existed 10 kya at the origins of adaptive radiation on San Salvador; furthermore, field fitness experiments demonstrate that the current environment continues to drive diversification in these lakes in the presence of ongoing gene flow (Martin and Wainwright 2013b; Martin and Feinstein 2014). Additional unmeasured ecological variables also appear comparable between San Salvador and neighboring saline lakes: all lakes are physiochemically similar with shared carbonate geology, the same dominant macroalgae species, and the same 1-2 insectivorous fish species (Godfrey 1994; Rothfus 2012). Primary productivity and pupfish population densities also appeared similar across islands based on observations of terrestrial inputs and catch per unit effort. The exceptional diversification rates observed on San Salvador relative to neighboring generalist populations would suggest the presence of exceptional ecological differences unlikely to be overlooked. Surprisingly, this survey indicates that any ecological differences unique to San Salvador are subtle, if present. Perhaps rare macroalgae taxa are necessary to facilitate additional fitness peaks by providing nutritional or structural substrate for critical species of gastropods or ostracods, the predominant food source of the molluscivore pupfish, or shelter for the scale-eating pupfish to successfully ambush its prey. Although alternative stable ecosystem states often exhibit sensitive dependence on initial ecological conditions (Beisner et al. 2003), such subtle thresholds and sensitivity would be unprecedented within the ecological theory of adaptive radiation (Simpson 1944; Schluter 2000; Gavrilets and Losos 2009; Erwin 2015b; but see Chen et al. 2015).

Conversely, the abundance of scales in all pupfish communities and pervasiveness of gene flow would suggest that scale-eating pupfish should colonize the entire Caribbean. There was no unique signature of secondary gene flow into San Salvador as predicted by the hybrid swarm hypothesis (Seehausen 2004); instead, all current evidence suggests that gene flow is ubiquitous and ongoing among islands and lakes (Fig. 5; Martin and Feinstein 2014). If sufficient ecological opportunity and genetic diversity exists in most Caribbean saline lakes, what are the additional constraints on adaptive diversification and dispersal of specialists across the Caribbean? One possibility is that colonization of neighboring populations may be constrained by the extremely low fitness of scale-eater hybrids (demonstrated in Martin and Wainwright 2013b; Martin 2016) and low probability of dispersal of a scale-eater pair to a neighboring lake due to their low frequency in all lake populations [<5% (Martin and Wainwright 2013c)]. Indeed, fitness experiments with F2 hybrid pupfish on San Salvador indicated that a large fitness valley separates scale-eating from generalist phenotypes, suggesting that survival in the wild is dependent on an extreme scale-eating phenotype with the full complement of adaptive alleles underlying this complex phenotype, whereas intermediate phenotypes reside in a deep fitness valley (Martin and Wainwright 2013a; Martin 2016; Martin et al. in review). Thus, the complex genetic architecture of the scale-eater phenotype and its interaction with fitness may further constrain its evolution, even in the presence of sufficient ecological opportunity.

Ultimately, this comprehensive examination of the origins of novel ecological niches during adaptive radiation rejects the ecological and hybrid swarm theories of adaptive radiation. The paradox is why 1000-fold differences in the diversification of trophic traits are not associated with 1000-fold differences in ecological opportunity or genetic diversity? Admittedly, the relationship between these two variables is unlikely to be one-to-one; however, it is worth emphasizing the dramatic nonlinearity between trait diversification rates and ecological or genetic explanatory variables within this system.

An increasing number of case studies suggests that the nonlinear emergence of novel traits leading to evolutionary innovations may be limited by the availability of genetic variation for complex phenotypes in addition to the presence of ecological opportunity (Schuettpelz and Pryer 2009; Wilson et al. 2012, Erwin 2015a; Harmon and Harrison 2015). For example, in the Lenski long-term evolution experiment, the evolution of citrate-feeding was limited by the specific mutations necessary for this major metabolic transition despite the continual presence of citrate (Blount et al. 2008, 2012; Quandt et al. 2015), providing a model for the rare origins of trophic novelty in pupfishes despite widespread examples of local adaptation throughout their range (e.g. Tobler and Carson 2010; Martin and Wainwright 2011). In contrast to parallel ecological speciation across similar environments, major ecological novelties, such as scale-eating, may emerge from non-ecological, contingent processes within a long-term background of ecological abundance. The isolation of novel ecological niches on the fitness landscape, as demonstrated for the scale-eating pupfish, may constrain their evolution even in the presence of resource abundance (e.g. scales in all pupfish communities), which could explain the singular evolution of scale-eating among thousands of Cyprinodontiform fishes: perhaps Caribbean pupfishes occupy a sparse fitness landscape dominated by a single wide peak (generalist algae-eating) with very few and difficult-to-access specialist peaks (e.g. scale-eating, molluscivory, piscivory, planktivory). Abundant ecological opportunity alone should not be expected to trigger such rare transitions.

Caribbean pupfishes provide a rare opportunity to examine the origins of evolutionary novelty. Such novelties are commonplace in many classic adaptive radiations (Martin and Wainwright 2013c), but can rarely be observed to vary across very similar populations in comparable neighboring environments. Despite Caribbean-wide gene flow and an abundance of competitor-free environments, trophic innovation is confined to a single island, providing a strong counterexample to the prevailing view of ecologically driven diversification and innovation.

## Acknowledgements

This study was funded by a Daphne and Ted Pengelley Award from the Center for Population Biology and ARCS Foundation awards to CHM. I thank the members of the American Killifish Association for supplying outgroup tissues, including B Turner, A Morales, F Vermuelen; R Hanna and K Guerrero for assistance obtaining permits; T Tran for measuring morphological specimens; Gerace Research Centre for accommodation; governments of the Bahamas and Dominican Republic for permission to conduct this research; and the Bahamian and Dominican people for their generous hospitality and assistance.

## References

Abbott, R., D. Albach, S. Ansell, J. W. Arntzen, S. J. E. Baird, N. Bierne, J. Boughman, A. Brelsford, C. A. Buerkle, R. Buggs, R. K. Butlin, U. Dieckmann, F. Eroukhmanoff, A. Grill, S. H. Cahan, J. S. Hermansen, G. Hewitt, A. Hudson, C. Jiggins, J. Jones, B. Keller, T. Marczewski, J. Mallet, P. Martinez-Rodriguez, M. Möst, S. Mullen, R. Nichols, A. W. Nolte, C. Parisod, K. Pfennig, A. M. Rice, M. G. Ritchie, B. Seifert, C. M. Smadja, R. Stelkens, J. M. Szymura, R. Väinölä, J. B. W. Wolf, and D. Zinner. 2013. Hybridization and speciation. J. Evol. Biol. 26:229–46.

Ackerly, D. 2009. Conservatism and diversification of plant functional traits: evolutionary rates versus phylogenetic signal. Proc. Natl. Acad. Sci. U. S. A. 106 Suppl:19699–706.

Alfaro, M. E., C. D. Brock, B. L. Banbury, and P. C. Wainwright. 2009. Does evolutionary innovation in pharyngeal jaws lead to rapid lineage diversification in labrid fishes? BMC Evol. Biol. 9:255.

Arbogast, B. S., S. V Drovetski, R. L. Curry, P. T. Boag, G. Seutin, P. R. Grant, B. R. Grant, and D. J. Anderson. 2006. The origin and diversification of Galapagos mockingbirds. Evolution 60:370–82.

Barkman, T. J., M. Bendiksby, S.-H. Lim, K. M. Salleh, J. Nais, D. Madulid, and T. Schumacher. 2008. Accelerated rates of floral evolution at the upper size limit for flowers. Curr. Biol. 18:1508–13.

Beisner, B., D. Haydon, and K. Cuddington. 2003. Alternative stable states in ecology. Front. Ecol. Environ. 1:376–382.

Berner, D., and W. Salzburger. 2015. The genomics of organismal diversification illuminated by adaptive radiations. Trends Genet. 31:491–499.

Blount, Z. D., J. E. Barrick, C. J. Davidson, and R. E. Lenski. 2012. Genomic analysis of a key innovation in an experimental *Escherichia coli* population. Nature 489:513–518.

Blount, Z. D., C. Z. Borland, and R. E. Lenski. 2008. Historical contingency and the evolution of a key innovation in an experimental population of *Escherichia coli*. Proc. Natl. Acad. Sci. U. S. A. 105:7899–906.

Bolnick, D. I., and O. L. Lau. 2008. Predictable patterns of disruptive selection in stickleback in postglacial lakes. Am. Nat. 172:1–11.

Bouckaert, R., J. Heled, D. Kühnert, T. Vaughan, C.-H. Wu, D. Xie, M. A. Suchard, A. Rambaut, and A. J. Drummond. 2014. BEAST 2: a software platform for bayesian evolutionary analysis. PLoS Comput. Biol. 10:e1003537.

Brawand, D., C. E. Wagner, Y. I. Li, M. Malinsky, I. Keller, S. Fan, O. Simakov, A. Y. Ng, Z. W. Lim, E. Bezault, J. Turner-Maier, J. Johnson, R. Alcazar, H. J. Noh, P. Russell, B. Aken, J. Alföldi, C. Amemiya, N. Azzouzi, J.-F. Baroiller, F. Barloy-Hubler, A. Berlin, R. Bloomquist, K. L. Carleton, M. a. Conte, H. D’Cotta, O. Eshel, L. Gaffney, F. Galibert, H. F. Gante, S. Gnerre, L. Greuter, R. Guyon, N. S. Haddad, W. Haerty, R. M. Harris, H. a. Hofmann, T. Hourlier, G. Hulata, D. B. Jaffe, M. Lara, A. P. Lee, I. MacCallum, S. Mwaiko, M. Nikaido, H. Nishihara, C. Ozouf-Costaz, D. J. Penman, D. Przybylski, M. Rakotomanga, S. C. P. Renn, F. J. Ribeiro, M. Ron, W. Salzburger, L. Sanchez-Pulido, M. E. Santos, S. Searle, T. Sharpe, R. Swofford, F. J. Tan, L. Williams, S. Young, S. Yin, N. Okada, T. D. Kocher, E. a. Miska, E. S. Lander, B. Venkatesh, R. D. Fernald, A. Meyer, C. P. Ponting, J. T. Streelman, K. Lindblad-Toh, O. Seehausen, and F. Di Palma. 2014. The genomic substrate for adaptive radiation in African cichlid fish. Nature 513:375–381.

Bryant, D., R. Bouckaert, J. Felsenstein, N. A. Rosenberg, and A. Roy Choudhury. 2012. Inferring species trees directly from biallelic genetic markers: bypassing gene trees in a full coalescent analysis. Mol. Biol. Evol. 29:1917–32.

Burnham, and Anderson. 2002. Model selection and multimodal inference: a practical information-theoretic approach. Springer Science and Business Media.

Burns, K. J., S. J. Hackett, and N. K. Klein. 2002. Phylogenetic relationships and morphological diversity in Darwin’s finches and their relatives. Evolution (N. Y). 56:1240.

Butler, M. A., and A. A. King. 2004. Phylogenetic comparative analysis: a modeling approach for adaptive evolution. Am. Nat. 164:683–695.

Catchen, J., P. A. Hohenlohe, S. Bassham, A. Amores, and W. A. Cresko. 2013. Stacks: an analysis tool set for population genomics. Mol. Ecol. 22:3124–40.

Catchen, J. M., A. Amores, P. Hohenlohe, W. Cresko, and J. H. Postlethwait. 2011. Stacks: building and genotyping loci de novo from short-read sequences. G3 1:171–82.

Chen, X., H.-F. Ling, D. Vance, G. a Shields-Zhou, M. Zhu, S. W. Poulton, L. M. Och, S.-Y. Jiang, D. Li, L. Cremonese, and C. Archer. 2015. Rise to modern levels of ocean oxygenation coincided with the Cambrian radiation of animals. Nat. Commun. 6:1–7.

Degnan, J. H., and N. Rosenberg. 2009. Gene tree discordance, phylogenetic inference and the multispecies coalescent. Trends Ecol. Evol. 24:332–40.

Degnan, J. H., and N. A. Rosenberg. 2006. Discordance of species trees with their most likely gene trees. PLoS Genet. 2:e68.

DeJong, T. M. 1975. A comparison of three diversity indices based on their components of richness and evenness. Oikos 26:222–227.

Dingerkus, G., and L. D. Uhler. 1977. Enzyme clearing of alcian blue stained whole small vertebrates for demonstration of cartilage. Stain Technol. 52:229–232.

Drummond, A. J., and A. Rambaut. 2007. BEAST: Bayesian evolutionary analysis by sampling trees. BMC Evol. Biol. 8:1–8.

Durand, E. Y., N. Patterson, D. Reich, and M. Slatkin. 2011. Testing for ancient admixture between closely related populations. Mol. Biol. Evol. 28:2239–52.

Eastman, J. M., M. E. Alfaro, P. Joyce, A. L. Hipp, and L. J. Harmon. 2011. A novel comparative method for identifying shifts in the rate of character evolution on trees. Evolution 65:3578–89.

Elena, S. F., and R. E. Lenski. 2003. Evolution experiments with microorganisms: the dynamics and genetic bases of adaptation. Nat. Rev. Genet. 4:457–69.

Elshire, R., J. Glaubitz, Q. Sun, J. Poland, K. Kawamoto, E. Buckler, and S. Mitchell. 2011. A robust, simple genotyping-by-sequencing (GBS) approach for high diversity species. PLoS One 6:e19379.

Erwin, D. H. 2015a. Novelty and innovation in the history of life. Curr. Biol. 25:R930–R940.

Erwin, D. H. 2015b. Was the Ediacaran–Cambrian radiation a unique evolutionary event? Paleobiology 41:1–15.

Evanno, G., S. Regnaut, and J. Goudet. 2005. Detecting the number of clusters of individuals using the software STRUCTURE: a simulation study. Mol. Ecol. 14:2611–20.

Felsenstein, J. 1985. Phylogenies and the comparative method. Am. Nat. 125:1–15.

Gavrilets, S., and J. B. Losos. 2009. Adaptive radiation: contrasting theory with data. Science 323:732–7.

Gillespie, R. 2004. Community assembly through adaptive radiation in Hawaiian spiders. Science 303:356–9.

Givnish, T. J., K. J. Sytsma, J. Smith, W. Hahn, B. DH, and E. Burkhardt. 1997. Molecular evolution and adaptive radiation in *Brocchinia* (Bromeliaceae: Pitcairnioideae) atop tepuis of the Guyana Shield. Pp. 259–311 in Molecular evolution and adaptive radiation. Cambridge University Press, Cambridge.

Godfrey, P. 1994. Natural history of northeastern San Salvador Island: A “New World” where the New World began. Bahamian Field Station.

Hansen, T. F. 1997. Stabilizing selection and the comparative analysis of adaptation: Evolution. 51:1341–1351.

Hansen, T. F., J. Pienaar, and S. H. Orzack. 2008. A comparative method for studying adaptation to a randomly evolving environment. Evolution 62:1965–77.

Harmon, L. J., and S. Harrison. 2015. Species diversity is dynamic and unbounded at local and continental scales. Am. Nat. 185:584–593.

Harmon, L. J., J. B. Losos, T. J. Davies, R. G. Gillespie, J. L. Gittleman, W. B. Jennings, K. H. Kozak, M. A. McPeek, F. Moreno-Roark, T. J. Near, A. Purvis, R. E. Ricklefs, D. Schluter, J. A. Schulte, O. Seehausen, B. L. Sidlauskas, O. Torres-Carvajal, J. T. Weir, and A. Ø. Mooers. 2010. Early bursts of body size and shape evolution are rare in comparative data. Evolution 64:2385–2396.

Harmon, L. J., J. T. Weir, C. D. Brock, R. E. Glor, and W. Challenger. 2008. GEIGER: investigating evolutionary radiations. Bioinformatics 24:129–31.

Heled, J., and A. J. Drummond. 2010. Bayesian inference of species trees from multilocus data. Mol. Biol. Evol. 27:570–80.

Horstkotte, J., and U. Strecker. 2005. Trophic differentiation in the phylogenetically young Cyprinodon species flock (Cyprinodontidae, Teleostei) from Laguna Chichancanab (Mexico). Biol. J. Linn. Soc. 85:125–134.

Hulsey, C. D., M. C. Mims, N. F. Parnell, and J. T. Streelman. 2010. Comparative rates of lower jaw diversification in cichlid adaptive radiations. J. Evol. Biol. 23:1456–1467.

Humphries, J. M. 1984. *Cyprinodon verecundus*, n. sp., a fifth species of pupfish from Laguna Chichancanab. Copeia 1984:58–68.

Humphries, J., and R. R. Miller. 1981. A remarkable species flock of pupfishes, genus *Cyprinodon*, from Yucatan, Mexico. Copeia 1981:52–64.

Hunter, J. P. 1998. Key innovations and the ecology of macroevolution. Trends Ecol. Evol. 13:31–6.

Janovetz, J. 2005. Functional morphology of feeding in the scale-eating specialist *Catoprion mento*. J. Exp. Biol. 208:4757–68.

Kassen, R., M. Llewellyn, and P. B. Rainey. 2004. Ecological constraints on diversification in a model adaptive radiation. Nature 431:984–988.

Kearse, M., R. Moir, A. Wilson, S. Stones-Havas, M. Cheung, S. Sturrock, S. Buxton, A. Cooper, S. Markowitz, C. Duran, T. Thierer, B. Ashton, P. Meintjes, and A. Drummond. 2012. Geneious Basic: an integrated and extendable desktop software platform for the organization and analysis of sequence data. Bioinformatics 28:1647–9.

Kisel, Y., and T. Barraclough. 2010. Speciation has a spatial scale that depends on levels of gene flow. Am. Nat. 175:316–334.

Kubatko, L. S., and J. H. Degnan. 2007. Inconsistency of phylogenetic estimates from concatenated data under coalescence. Syst. Biol. 56:17–24.

Kuhlwilm, M., I. Gronau, M. J. Hubisz, C. de Filippo, J. Prado-Martinez, M. Kircher, Q. Fu, H. A. Burbano, C. Lalueza-Fox, M. de la Rasilla, A. Rosas, P. Rudan, D. Brajkovic, Ž. Kucan, I. Gušic, T. Marques-Bonet, A. M. Andrés, B. Viola, S. Pääbo, M. Meyer, A. Siepel, and S. Castellano. 2016. Ancient gene flow from early modern humans into Eastern Neanderthals. Nature 530:429–433.

Lamichhaney, S., J. Berglund, M. S. Almén, K. Maqbool, M. Grabherr, A. Martinez-Barrio, M. Promerová, C.-J. Rubin, C. Wang, N. Zamani, B. R. Grant, P. R. Grant, M. T. Webster, and L. Andersson. 2015. Evolution of Darwin’s finches and their beaks revealed by genome sequencing. Nature 518:371–375.

Langerhans, R. B. 2010. Predicting evolution with generalized models of divergent selection: a case study with Poeciliid fish. Integr. Comp. Biol. 50:1167–1184.

Langerhans, R. B., M. E. Gifford, and E. O. Joseph. 2007. Ecological speciation in Gambusia fishes. Evolution 61:2056–74.

Langmead, B., and S. Salzberg. 2012. Fast gapped-read alignment with Bowtie 2. Nat. Methods 9:357–359.

Layman, C. A. 2007. What can stable isotope ratios reveal about mangroves as fish habitat? Bull. Mar. Sci. 80:513–527.

Lazaridis, I., N. Patterson, A. Mittnik, G. Renaud, S. Mallick, K. Kirsanow, P. H. Sudmant, J. G. Schraiber, S. Castellano, M. Lipson, B. Berger, C. Economou, R. Bollongino, Q. Fu, K. I. Bos, S. Nordenfelt, H. Li, C. de Filippo, K. Prüfer, S. Sawyer, C. Posth, W. Haak, F. Hallgren, E. Fornander, N. Rohland, D. Delsate, M. Francken, J.-M. Guinet, J. Wahl, G. Ayodo, H. a. Babiker, G. Bailliet, E. Balanovska, O. Balanovsky, R. Barrantes, G. Bedoya, H. Ben-Ami, J. Bene, F. Berrada, C. M. Bravi, F. Brisighelli, G. B. J. Busby, F. Cali, M. Churnosov, D. E. C. Cole, D. Corach, L. Damba, G. van Driem, S. Dryomov, J.-M. Dugoujon, S. a. Fedorova, I. Gallego Romero, M. Gubina, M. Hammer, B. M. Henn, T. Hervig, U. Hodoglugil, A. R. Jha, S. Karachanak-Yankova, R. Khusainova, E. Khusnutdinova, R. Kittles, T. Kivisild, W. Klitz, V. Kučinskas, A. Kushniarevich, L. Laredj, S. Litvinov, T. Loukidis, R. W. Mahley, B. Melegh, E. Metspalu, J. Molina, J. Mountain, K. Näkkäläjärvi, D. Nesheva, T. Nyambo, L. Osipova, J. Parik, F. Platonov, O. Posukh, V. Romano, F. Rothhammer, I. Rudan, R. Ruizbakiev, H. Sahakyan, A. Sajantila, A. Salas, E. B. Starikovskaya, A. Tarekegn, D. Toncheva, S. Turdikulova, I. Uktveryte, O. Utevska, R. Vasquez, M. Villena, M. Voevoda, C. a. Winkler, L. Yepiskoposyan, P. Zalloua, T. Zemunik, A. Cooper, C. Capelli, M. G. Thomas, A. Ruiz-Linares, S. a. Tishkoff, L. Singh, K. Thangaraj, R. Villems, D. Comas, R. Sukernik, M. Metspalu, M. Meyer, E. E. Eichler, J. Burger, M. Slatkin, S. Pääbo, J. Kelso, D. Reich, and J. Krause. 2014. Ancient human genomes suggest three ancestral populations for present-day Europeans. Nature 513:409–413.

Lencer, E. S., M. L. Riccio, and A. R. McCune. 2016. Changes in growth rates of oral jaw elements produce evolutionary novelty in Bahamian pupfish. J. Morphol. in press.

Liem, K. 1980. Acquisition of energy by teleosts: adaptive mechanisms and evolutionary patterns. Pp. 299–334 *in* Environmental Physiology of Fishes.

Liu, L., Z. Xi, S. Wu, C. C. Davis, and S. V. Edwards. 2015. Estimating phylogenetic trees from genome-scale data. Ann. N. Y. Acad. Sci. 1360:36–53.

Losos, J. 2009. Lizards in an evolutionary tree: ecology and adaptive radiation of anoles. University of California Press, Berkeley.

Losos, J. B., and D. Schluter. 2000. Analysis of an evolutionary species-area relationship. Nature 408:847–50.

Losos, J., and D. Mahler. 2010. Adaptive radiation: the interaction of ecological opportunity, adaptation, and speciation. Pp. 381–420 *in* Evolution Since Darwin: the first 150 years. Sinauer Associates, Sunderland, MA.

Malinsky, A. M. T. Richard J. Challis, S. Schiffels, Y. Terai, B. P. Ngatunga, E. A. Miska, R. Durbin, M. J. Genner, and G. F. Turner. 2015. Genomic islands of speciation separate cichlid ecomorphs in an East African crater lake. Science (80-.). 350:1493–1498.

Martin, C. 2016. Context-dependence in complex adaptive landscapes: frequency and trait-dependent selection surfaces within an adaptive radiation of Caribbean pupfishes. Evolution (N. Y). DOI: 10.1111/evo.12932.

Martin, C. 2013. Strong assortative mating by diet, color, size, and morphology but limited progress toward sympatric speciation in a classic example: Cameroon crater lake cichlids. Evolution 67:2114–23.

Martin, C. 2012. Weak disruptive selection and incomplete phenotypic divergence in two classic examples of sympatric speciation: cameroon crater lake cichlids. Am. Nat. 180:E90–E109.

Martin, C. H., J. E. Crawford, B. J. Turner, L. H. Simons, and C. H. Martin. 2016. Diabolical survival in Death Valley: recent pupfish colonization, gene flow, and genetic assimilation in the smallest species range on earth. Proc. R. Soc. B Biol. Sci. 283:23–34.

Martin, C. H., J. S. Cutler, J. P. Friel, T. Dening, G. Coop, and P. C. Wainwright. 2015. Complex histories of repeated colonization and hybridization cast doubt on the clearest examples of sympatric speciation in the wild. Evolution (N. Y). 69:1406–1422.

Martin, C. H., and L. C. Feinstein. 2014. Novel trophic niches drive variable progress towards ecological speciation within an adaptive radiation of pupfishes. Mol. Ecol. 23:1846–62.

Martin, C. H., and M. J. Genner. 2009. High niche overlap between two successfully coexisting pairs of Lake Malawi cichlid fishes. Can. J. Fish. Aquat. Sci. 66:579–588.

Martin, C. H., and P. C. Wainwright. 2013a. A remarkable species flock of Cyprinodon pupfishes endemic to San Salvador Island, Bahamas. Bull. Peabody Museum Nat. Hist. 54:231–240.

Martin, C. H., and P. C. Wainwright. 2013b. Multiple fitness peaks on the adaptive landscape drive adaptive radiation in the wild. Science 339:208–211.

Martin, C. H., and P. C. Wainwright. 2013c. On the measurement of ecological novelty: scale-eating pupfish are separated by 168 my from other scale-eating fishes. PLoS One 8:e71164.

Martin, C. H., and P. C. Wainwright. 2011. Trophic novelty is linked to exceptional rates of morphological diversification in two adaptive radiations of Cyprinodon pupfishes. Evolution 65:2197–212.

Martins, E. P., and T. F. Hansen. 1997. Phylogenies and the comparative method: A general approach to incorporating phylogenetic information into the analysis of interspecific data. Am. Nat. 149:646.

McKinnon, J. S., S. Mori, B. K. Blackman, L. David, D. M. Kingsley, L. Jamieson, J. Chou, and D. Schluter. 2004. Evidence for ecology’s role in speciation. Nature 429:294–298.

McVean, G. 2009. A genealogical interpretation of principal components analysis. PLoS Genet. 5:e1000686.

Miller, M. A., W. Pfeiffer, and T. Schwartz. 2010. Creating the CIPRES Science Gateway for inference of large phylogenetic trees. 2010 Gatew. Comput. Environ. Work. 1–8. Ieee.

Nosil. 2012. Ecological speciation. Oxford University Press, Oxford.

Nosil, P., L. J. Harmon, and O. Seehausen. 2009. Ecological explanations for (incomplete) speciation. Trends Ecol. Evol. 24:145–56. Elsevier Ltd.

O’Meara, B. C., C. Ané, M. J. Sanderson, and P. C. Wainwright. 2006. Testing for different rates of continuous trait evolution using likelihood. Evolution 60:922–33.

Parent, C. E., and B. J. Crespi. 2009. Ecological opportunity in adaptive radiation of Galápagos endemic land snails. Am. Nat. 174:898–905.

Pfennig, D. W., and K. S. Pfennig. 2010. Character displacement and the origins of diversity. Am. Nat. 176 Suppl:S26–44.

Pfennig, D. W., and K. S. Pfennig. 2012. Evolution’s wedge: competition and the origins of diversity. University of California Press.

Pickrell, J. K., and J. K. Pritchard. 2012. Inference of population splits and mixtures from genome-wide allele frequency data. PLoS Genet. 8:e1002967.

Post, D. 2002. Using stable isotopes to estimate trophic position: models, methods, and assumptions. Ecology 83:703–718.

Price, S. a, R. Holzman, T. J. Near, and P. C. Wainwright. 2011. Coral reefs promote the evolution of morphological diversity and ecological novelty in labrid fishes. Ecol. Lett. 14:462–9.

Purcell, S., B. Neale, and K. Todd-Brown. 2007. PLINK: a tool set for whole-genome association and population-based linkage analyses. Am. J. Hum. Genet. 81:559–575.

Puritz, J. B., M. V Matz, R. J. Toonen, J. N. Weber, D. I. Bolnick, and C. E. Bird. 2014. Demystifying the RAD fad. Mol. Ecol. 23:5937–42.

Quandt, E. M., J. Gollihar, Z. D. Blount, A. D. Ellington, G. Georgiou, and J. E. Barrick. 2015. Fine-tuning citrate synthase flux potentiates and refines metabolic innovation in the Lenski evolution experiment. Elife 4:e09696.

R Core Team. 2015. R: A language and environment for statistical computing. R Foundation for Statistical Computing, Vienna, Austria.

Rainey, P., and M. Travisano. 1998. Adaptive radiation in a heterogeneous environment. Nature 394:69–72.

Reich, D., K. Thangaraj, N. Patterson, A. L. Price, and L. Singh. 2009. Reconstructing Indian population history. Nature 461:489–94.

Revell, L. J. 2009. Size-correction and principal components for interspecific comparative studies. Evolution 63:3258–68.

Rieseberg, L. H., O. Raymond, D. M. Rosenthal, Z. Lai, K. Livingstone, T. Nakazato, J. L. Durphy, A. E. Schwarzbach, L. A. Donovan, and C. Lexer. 2003. Major ecological transitions in wild sunflowers facilitated by hybridization. Science 301:1211–6.

Roderick, G. K., and R. G. Gillespie. 1998. Speciation and phylogeography of Hawaiian terrestrial arthropods. Mol. Ecol. 7:519–31.

Rohlf, F. 2001. TPSDig2: a program for landmark development and analysis.

Rothfus, E. 2012. Water-quality monitoring of San Salvadorian inland lakes. Pp. 129–138 *in* D. Gamble and P. Kindler, eds. Proceedings of the 15th Symposium on the Geology of the Bahamas and other Carbonate Regions. Gerace Research Center.

Roy, D., K. Lucek, R. P. Walter, and O. Seehausen. 2015. Hybrid “superswarm” leads to rapid divergence and establishment of populations during a biological invasion. Mol. Ecol. 24:5394–5411.

Rundle, H. D., and P. Nosil. 2005. Ecological speciation. Ecol. Lett. 8:336–352.

Santini, F., M. T. T. Nguyen, L. Sorenson, T. B. Waltzek, J. W. Lynch Alfaro, J. M. Eastman, and M. E. Alfaro. 2013. Do habitat shifts drive diversification in teleost fishes? An example from the pufferfishes (Tetraodontidae). J. Evol. Biol. 26:1003–1018.

Sazima, I. 1983. Scale-eating in characoids and other fishes. Environ. Biol. Fishes 9:87–101.

Schluter, D. 2000. Ecology of adaptive radiation. Oxford University Press, Oxford.

Schmitter-Soto, J. J. 1999. Distribution of continental fishes in northern Quintana Roo, Mexico. Southwest. Nat. 44:166–172.

Schuettpelz, E., and K. Pryer. 2009. Evidence for a Cenozoic radiation of ferns in an angiosperm-dominated canopy. Proc. Natl. Acad. Sci. U. S. A. 106:11200–11205.

Schumer, M., R. Cui, B. Boussau, R. Walter, G. Rosenthal, and P. Andolfatto. 2013. An evaluation of the hybrid speciation hypothesis for Xiphophorus clemenciae based on whole genome sequences. Evolution (N. Y). 67:1155–1168.

Schumer, M., R. Cui, G. G. Rosenthal, and P. Andolfatto. 2015. Reproductive isolation of hybrid populations driven by genetic incompatibilities. PLoS Genet. 11:e1005041.

Seehausen, O. 2006. African cichlid fish: a model system in adaptive radiation research. Proc. Biol. Sci. 273:1987–98.

Seehausen, O. 2004. Hybridization and adaptive radiation. Trends Ecol. Evol. (Personal Ed. 19:198–207.

Seehausen, O., R. K. Butlin, I. Keller, C. E. Wagner, J. W. Boughman, P. A. Hohenlohe, C. L. Peichel, G.-P. Saetre, C. Bank, A. Brännström, A. Brelsford, C. S. Clarkson, F. Eroukhmanoff, J. L. Feder, M. C. Fischer, A. D. Foote, P. Franchini, C. D. Jiggins, F. C. Jones, A. K. Lindholm, K. Lucek, M. E. Maan, D. A. Marques, S. H. Martin, B. Matthews, J. I. Meier, M. Möst, M. W. Nachman, E. Nonaka, D. J. Rennison, J. Schwarzer, E. T. Watson, A. M. Westram, and A. Widmer. 2014. Genomics and the origin of species. Nat. Rev. Genet. 15:176–92.

Servedio, M. R., J. Hermisson, and G. S. van Doorn. 2013. Hybridization may rarely promote speciation. J. Evol. Biol. 26:282–5.

Simpson, G. G. 1944. Tempo and mode of evolution. 15th ed. Columbia University Press.

Stacklies, W., H. Redestig, M. Scholz, D. Walther, and J. Selbig. 2007. pcaMethods–a bioconductor package providing PCA methods for incomplete data. Bioinformatics 23:1164–7.

Stankowski, S., and M. A. Streisfeld. 2015. Introgressive hybridization facilitates adaptive divergence in a recent radiation of monkeyflowers. Proc. R. Soc. London B 282:20151666.

Stevenson, M. M. 1992. Food habits within the Laguna Chichancanab *Cyprinodon* (Pisces: Cyprinodontidae) species flock. Southwest. Nat. 37:337–343.

Strecker, U. 2006a. Genetic differentiation and reproductive isolation in a Cyprinodon fish species flock from Laguna Chichancanab, Mexico. Mol. Phylogenet. Evol. 39:865–72.

Strecker, U. 2006b. The impact of invasive fish on an endemic Cyprinodon species flock (Teleostei) from Laguna Chichancanab, Yucatan, Mexico. Ecol. Freshw. Fish 15:408–418.

Tobler, M., and E. W. Carson. 2010. Environmental variation, hybridization, and phenotypic diversification in Cuatro Ciénegas pupfishes. J. Evol. Biol. 23:1475–89.

Uyeda, J. C., and L. J. Harmon. 2014. A novel Bayesian method for inferring and interpreting the dynamics of adaptive landscapes from phylogenetic comparative data. Syst. Biol. 63:902–918.

Vamosi, S. M. 2003. The presence of other fish species affects speciation in threespine sticklebacks. Evol. Ecol. Res. 5:717–730.

Venables, W., and B. Ripley. 2013. Modern applied statistics. Springer Science and Business Media.

Vila, I., P. Morales, S. Scott, E. Poulin, D. Véliz, C. Harrod, and M. a Méndez. 2013. Phylogenetic and phylogeographic analysis of the genus Orestias (Teleostei: Cyprinodontidae) in the southern Chilean Altiplano: the relevance of ancient and recent divergence processes in speciation. J. Fish Biol. 82:927–43.

Wagner, C. E., L. J. Harmon, and O. Seehausen. 2014. Cichlid species-area relationships are shaped by adaptive radiations that scale with area. Ecol. Lett. 17:583–92.

Wagner, C. E., L. J. Harmon, and O. Seehausen. 2012. Ecological opportunity and sexual selection together predict adaptive radiation. Nature 487:366–9.

Weinreich, D. M., and L. Chao. 2005. Rapid evolutionary escape by large populations from local fitness peaks is likely in nature. Evolution 59:1175–82.

Whittall, J. B., and S. A. Hodges. 2007. Pollinator shifts drive increasingly long nectar spurs in columbine flowers. Nature 447:706–9.

Wilson, G. P., a. R. Evans, I. J. Corfe, P. D. Smits, M. Fortelius, and J. Jernvall. 2012. Adaptive radiation of multituberculate mammals before the extinction of dinosaurs. Nature 483:457–460. Nature Publishing Group.

